# Lateral membrane organization as target of an antimicrobial peptidomimetic compound

**DOI:** 10.1101/2023.01.17.524350

**Authors:** Adéla Melcrová, Sourav Maity, Josef Melcr, Niels A. W. de Kok, Mariella Gabler, Jonne van der Eyden, Wenche Stensen, John S. M. Svendsen, Arnold J. M. Driessen, Siewert J. Marrink, Wouter H. Roos

**Affiliations:** Molecular Biophysics, Zernike institute for Advanced Materials, Rijksuniversiteit Groningen, the Netherlands; Molecular Dynamics, Groningen Biomolecular Sciences & Biotechnology Institute, Rijksuniversiteit Groningen, the Netherlands; Molecular Microbiology, Groningen Biomolecular Sciences & Biotechnology Institute, Rijksuniversiteit Groningen, the Netherlands; Department of Chemistry, UiT Arctic University of Norway, Norway

**Keywords:** Staphylococcus aureus, peptidomimetic, atomic force microscopy, MD simulations, bacterial membrane, lipid domains, antimicrobial resistance, lipidomics

## Abstract

Antimicrobial resistance is one of the leading concerns in medical care. Here we resolve the functional mechanism of the antimicrobial action of the cationic tripeptide AMC-109 by combining high speed-atomic force microscopy, molecular dynamics, fluorescence assays, and lipidomic analysis. We show that AMC-109 activity on the negatively charged plasma membrane of *Staphylococcus aureus* consists of two crucial steps. First, AMC-109 self-assembles into stable aggregates with specificity for negatively charged membranes. Second, by incorporation into the *S. aureus* membrane the lateral membrane organization is affected, dissolving membrane nanodomains. Domain dissolution affects membrane functions such as protein sorting and cell wall synthesis, and is suggested to cause a loss of resistance of methicillin-resistant *S. aureus* (MRSA) to methicillin. As the AMC-109 mode of action is similar to the activity of the disinfectant benzalkonium chloride (BAK), a broad applicability, but with low cytotoxicity to human cells, is expected.

## Introduction

Ever since the introduction of antibiotics, bacteria have developed resistance against these compounds. For instance, methicillin-resistant variants of the Gram-positive bacterium *Staphylococcus aureus* (MRSA) appeared already several decades ago^1^. This discovery spurred the search for new types of antibiotics, however, this quest has met only limited success. In fact, antimicrobial resistance was declared by the World Health Organization as one of the top 10 global threats in medical care, with *S aureus* as one of the leading pathogens^2^. Natural antimicrobial peptides and their mimics are promising scaffolds for the development of new antibiotics. Antimicrobial peptides are an integral part of our immune system, and as such they have potent antimicrobial activity against a broad range of bacteria^3–5^. However, they have several drawbacks, such as high production costs, cytotoxicity to red blood cells, and development of bacterial resistance^6,7^. One way to overcome the antimicrobial resistance is to stabilize the antimicrobial peptides by using non-native amino acids or modified peptide chains generating non-natural peptide mimics — peptidomimetics — which are stable in vivo, not harmful to human cells, and not suffering from resistance^5,7–10^. Here we focus on a promising peptidomimetic AMC-109 (Figure 1a)^11^, a cationic artificial tripeptide that is relatively simple to synthesize, has a broad antimicrobial activity, and can be processed in large-scale industry^7^. AMC-109 follows the minimal pharmacophore model developed in the lab of Svendsen^12^ stating a simple rule that a minimum of four amino acid residues are required for significant antibacterial activity — two of them cationic, the other two hydrophobic^7,11,12^. In AMC-109 the two hydrophobic residues are replaced by a single artificial residue encompassing the bulk and hydrophobicity of two individual tryptophans. Two arginine-like residues than provide the positive charge.

**Fig. 1:**
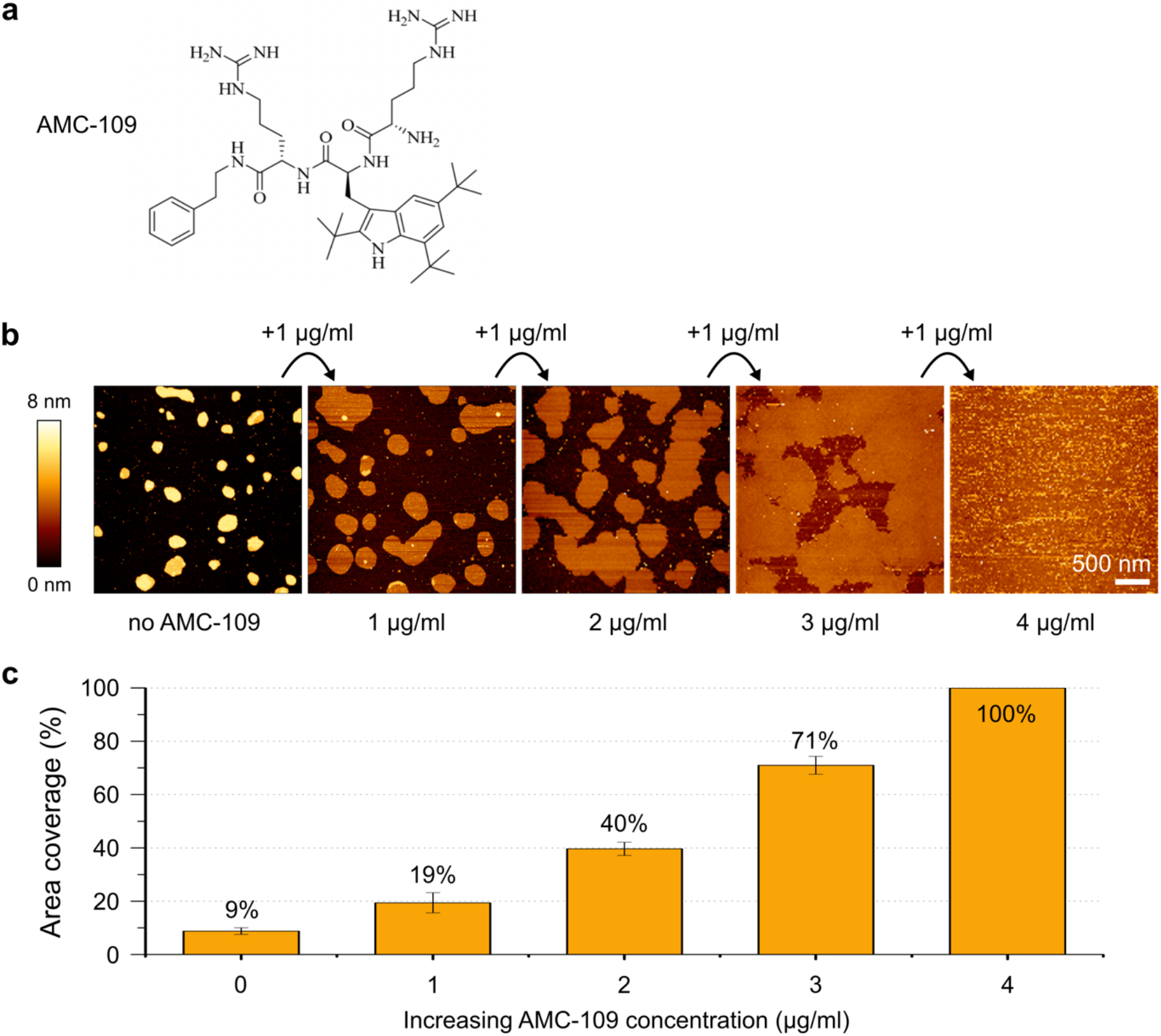
AFM imaging experiments reveal the effect of AMC-109 addition to *S. aureus* lipid membranes. **a** AMC-109 chemical structure. **b***S. aureus* lipid membranes upon treatment with AMC-109. Membranes are visualized untreated (left) and treated by subsequent additions of 1 μg/ml AMC-109 (0, 1, 2, 3, 4 μg/ml). **c** Area of the mica surface covered by the membrane upon subsequent additions of 1 μg/ml AMC-109. For each concentration, coverage of 10–12 areas in a 3×3 μm image size gathered from three independent experiments (representative example at **b**) were evaluated. Error bars represent standard error of the mean.

The antimicrobial activity of AMC-109 was positively tested against two Gram-negative bacteria, *Escherichia coli* and *Pseudomonas aeruginosa*, as well as a multitude of Gram-positive *S. aureus* strains including the strains resistant to common antibiotics such as methicillin or vancomycin^11,13^. Clinical trials bring promising results so far. In nasal decolonization studies conducted in Sweden, patients infected with two variants of *S. aureus* were treated for three days with AMC-109, which significantly reduced infection^14^. Integrating of AMC-109 into wound dressings and gels leads to significantly higher antibacterial activity compared to current clinical standard therapy^15^. Moreover, AMC-109 was demonstrated to have more than five times longer post-antibiotic effect against various *S. aureus* strains than commonly used mupirocin, and could hence be applied less frequently or in smaller doses to maintain its antimicrobial activity^16^. The promising pharmaceutical studies raise the question what the molecular mechanism of the antimicrobial activity of AMC-109 is. Detailed understanding of the antibiotic effects at the molecular level will support identifying new drug targets in resistant bacteria, and can be used in further development of new potent antimicrobial drugs. Here we test how AMC-109 affects the membranes of *S. aureus* on a molecular/nanoscopic level by a combination of atomic force microscopy (AFM), high-speed AFM (HS-AFM), fluorescence leakage assays, molecular dynamics (MD) simulations, mass spectrometry-based lipidomic analysis and gel chromatography, yielding a comprehensive view on the complex molecular mechanism of AMC-109 activity on the membrane of *S. aureus*.

## Results and discussion

### Lipid extracts as a model of *S. aureus* cytoplasmic membrane

In order to scrutinize the interactions of AMC-109 with the membrane, we extracted lipids from *S. aureus* cells^17^ to be used as bacterial model membranes for AFM and HS-AFM investigation. The lipids were characterized through lipidomic analyses (Table 1, Figure S1). The membrane of the *S. aureus* comprises of glycerophospholipids, followed by glycolipids^18–20^. Mass spectrometry and thin layer chromatography demonstrate that the most abundant glycerophospholipid is the negatively charged phosphatidylglycerol (PG, ~66% of all glycerophospholipids) and the most abundant glycolipid is diglycosyldiacylglycerol (~85% of all glycolipids). Overall, the lipidomic analysis confirms the presence of the negatively charged molecules in the phospholipid membranes of *S. aureus*, which results in the high attractivity for the cationic AMC-109. The analysis of the acyl chains distribution in the phospholipids fraction showed that the majority of the phospholipids contains saturated acyl chains, often with iso- or anteiso- branching complying with previous *S. aureus* lipidomic studies^18,21^. Moreover, the lipid extracts also contain a significant portion of unsaturated lipids (~10% of PG lipids, Figure S1f), which is typically not reported for *S. aureus* ^19,22^.

**Table 1.** Lipidomic analysis of total lipid extracts from *S. aureus* (Figure S1).

| Compound<sup>1</sup> | Charge at pH ~ 7 | Relative densitometric response (Molybdenum blue) |
| --- | --- | --- |
| CL | -2 | 9 % |
| PG | -1 | 66 % |
| Lysyl-PG | +1 | 24 % |
| Compound | | Relative densitometric response (α-naphthol) |
| 1-hex-DAG | 0 | 15 % |
| 2-hex-DAG | 0 | 85 % |

<sup>1</sup> CL, cardiolipin; PG, phosphatidylglycerol; Lysyl-PG, lysyl-phosphatidylglycerol; 1-hex-DAG, monoglycosyldiacylglycerol; 2-hex-DAG, diglycosyldiacylglycerol

### AMC-109 changes structural properties of *S. aureus* lipid membranes

AFM imaging was employed to observe the effects of AMC-109 on supported membranes prepared from the *S. aureus* lipid extracts. The AFM images at increasing concentrations of AMC-109 (Figure 1b) reveal effects on the membranes already at concentrations of 1–2 μg/ml. Interestingly, this concentration coincides well with the reported minimal inhibitory concentration (MIC) that induces antimicrobial activity on living *S. aureus*^11^. Upon exposure of the *S. aureus* lipid membranes to AMC-109 the membrane expands (Figure 1b). For concentrations up to 2 μg/ml, this is a relatively fast process reaching an equilibrium state after several minutes (Figure S2a). The expansion at 3 μg/ml is more gradual, slightly changing even after 40 minutes (Figure S2b). The relative surface coverage starts around 10% without AMC-109 and gradually increases to reach 100% at 4 μg/ml of the peptidomimetic (Figure 1b and c).

### AMC-109 clusters and dissolves lateral lipid domains in *S. aureus* lipid membranes

AFM imaging reveals a co-existence of regions with higher and lower thickness in the *S. aureus* lipid membranes (Figure S3). However, these regions were typically smeared out suggesting a dynamic nature of these regions. We, hence, moved to HS-AFM to improve the temporal resolution^23,24^. HS-AFM reveals that the higher thickness regions in the untreated *S. aureus* lipid membranes are highly mobile lateral lipid domains (Figure 2a) that maintain roughly circular shape with a strikingly narrow size distribution of ~30 nm in diameter (30.9 ± 0.4 nm, N = 269, Figure S4).

**Fig. 2:**
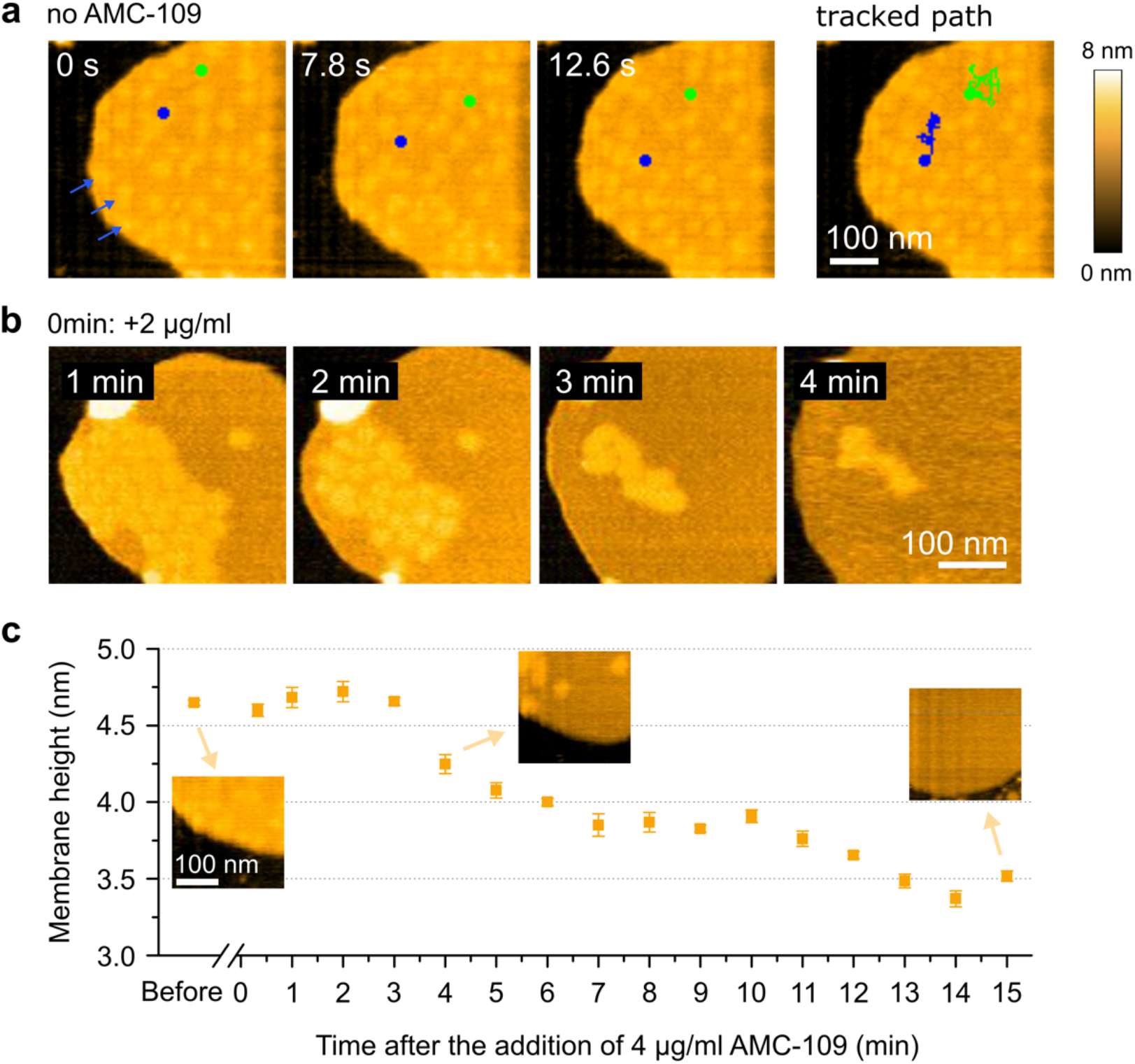
HS-AFM imaging of the AMC-109 effects on *S. aureus* lipid membranes supported on mica. **a** HS-AFM images of the untreated membrane showing the diffusion dynamics of lateral lipid domains (blue arrows). Dynamic path of two lateral domains (blue and green discs) were tracked in the 0.3 s per frame movie. **b** HS-AFM images of the membrane in time after the addition of 2 μg/ml AMC-109. For an example of a 0 min frame see panel **a**. Lateral domains accumulate together, then gradually dissolve. This is followed by expansion and thinning of the membrane. **c** Membrane thickness evolution over time after the addition of 4 μg/ml AMC-109. Measured from HS-AFM images at the continuous parts of the *S. aureus* lipid membrane surrounding the lateral lipid domains. Mean membrane height and standard errors of the mean are displayed. Data points are each the mean of N > 20 individual measurements (N = 363, 49, 22, 21, 74, 31, 52, 93, 37, 41, 71, 27, 35, 34, 27, 24, and 45, from left to right respectively). Insets shows membrane details before the AMC-109 addition, and at t = 4 min and t = 15 min after. Average height before the AMC-109 addition is 4.65 ± 0.02 nm. At time 13–15 min the height settles around 3.5 nm.

The observed phase separation probably arises from the complex composition of the lipid extracts, which apart from phospholipids also contain glycolipids and a portion of cardiolipin^25–28^. The domains display a rapid lateral movement inside the untreated membrane (Figure 2a). After the addition of AMC-109 at the concentrations above the MIC (Figure 2b), however, the domains accumulate together (Figure 2b 1–2 min) before they gradually disappear (Figure 2b 3–4 min). The accumulation of the domains occurs within tens of seconds after the addition of AMC-109 to the membranes. The accumulated domains gradually dissolve after about 3–5 minutes and only then the membrane starts expanding and significantly thinning (Figure 2b–c). Interestingly, lower AMC-109 concentrations (around 1 μg/ml) are enough to induce the accumulation and domain dissolution, which significantly alters the mechanical properties of the membrane, increasing its Young’s modulus (Figure S5). These concentrations, however, do not lead to full membrane expansion, which we observe only at higher AMC-109 concentrations. We measured the height of the *S. aureus* lipid membrane over time after adding 4 μg/ml AMC-109. The time evolution of the membrane height *h* is displayed in Figure 2c. A substantial thinning of ~1 nm of the continuous part of the membrane, i.e. the part not including the domains, is observed. In these experiments it is assured that the mica background does not get covered with AMC-109, allowing for accurate height measurements (Figure S5).

The disc-shaped lateral domains in our *S. aureus* lipid membranes (Figure 2a) are reminiscent of native functional membrane microdomains (FMM) from living bacteria^29^. FMMs in the cytoplasmic membrane of *S. aureus* fulfill important functions such as peptidoglycan synthesis, membrane lipid metabolism, membrane transport, protein quality control, virulence, and also oligomerization of low-affinity penicillin-binding protein (PBP2a) responsible for the resistance against penicillin, methicillin and other β-lactam antibiotics^30–34^. It was shown that FMMs contain cardiolipins and glycolipids^28,34^, which are also present in our *S. aureus* lipid membranes. Dissolution of FMMs in living bacteria leads to their death as the bacteria lose the multiple functions associated with these domains. Moreover, MRSA bacteria without the FMMs are no longer resistant to β-lactam antibiotics including methicillin^30,31,34^. We, hence, suggest that the AMC-109 treatment could be further enhanced by the use of β-lactam antibiotics.

### Molecular picture of AMC-109 attacking model membranes

We have employed MD simulations using the coarse grained Martini 3 force field^35,36^ to obtain a molecular picture of AMC-109 activity at model lipid membranes. Figure 3a shows the chemical structure of the AMC-109 molecule together with its Martini 3 representation and dimensions. The amphiphilic nature of AMC-109 molecules drives their fast spontaneous self-assembly into micellar aggregates within the first 100 ns of equilibration simulations^37^. Such spontaneously formed aggregates adopt a globular or slightly elongated shape with a characteristic size of 4 nm in diameter (Figure 3b).

**Fig. 3:**
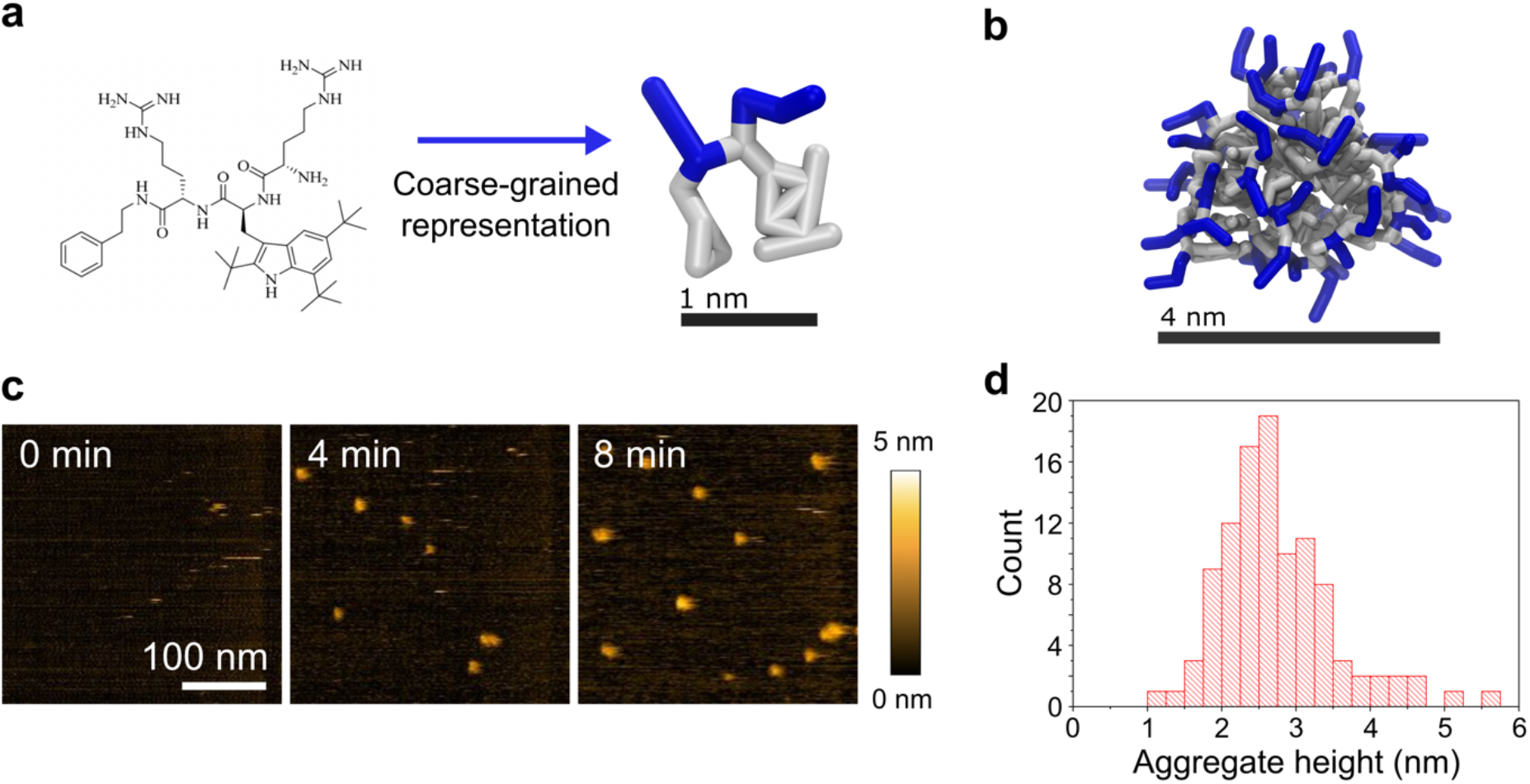
Aggregation properties of AMC-109. **a** AMC-109 chemical structure and its representation in Martini 3 coarse grained model. Cationic arginine-like residues are depicted in blue, artificial hydrophobic residue that encompass the hydrophobicity of two tryptophans (small triangle and the large planar structure) in grey. **b** Representative self-assembled aggregate of AMC-109 molecules formed after ~0.5 μs of the MD simulation. **c** HS-AFM images of micellar aggregates adsorbed on mica after the addition of 5 μg/ml AMC-109. **d** Histogram of the measured aggregate height. Average aggregate height was evaluated as 2.75 ± 0.08 nm (N = 104).

From both AFM and HS-AFM imaging, we see that such aggregates also attach to the mica surface. Figure 3c shows how after adding 5 μg/ml AMC-109 to the buffer solution, individual ball shaped aggregates appear on the mica. The height of these aggregates attached on a solid surface is 2.75 ± 0.08 nm (N = 104, Figure 3d) being in line with the size of a single aggregate in the bulk predicted by MD simulations (Figure 3b). After the attachment of the first aggregate on mica during our HS-AFM experiment, the adsorption process continues and the aggregates eventually form a continuous layer of molecules with a thickness of 1–3 nm.

Bacterial membranes possess a net negative charge (~70% negatively charged lipids in *S. aureus* lipid membranes^38^), while the outer leaflet of eukaryotic membranes is neutral^39,40^. To demonstrate the selectivity of AMC-109 to bacterial membranes, we have varied the surface charge of the simulated model membranes. The resulting series of MD simulations with mixed POPC/POPG bilayers (POPG content 0–100 mol%; simulation setup see Figure S8) demonstrates the increasing attractivity of the AMC-109 aggregates to membranes with a higher negative charge (Figure 4a). Notably, the adsorption of AMC-109 always saturates close to the point when the net charge of the membrane is neutralized. In particular, membranes with 20 mol% POPG or more adsorb on average one AMC-109 molecule (charge +3) per three molecules of POPG (charge −1), while membranes with small negative charge (≤ 10 mol% POPG) attract negligible amounts of AMC-109 molecules

**Fig 4:**
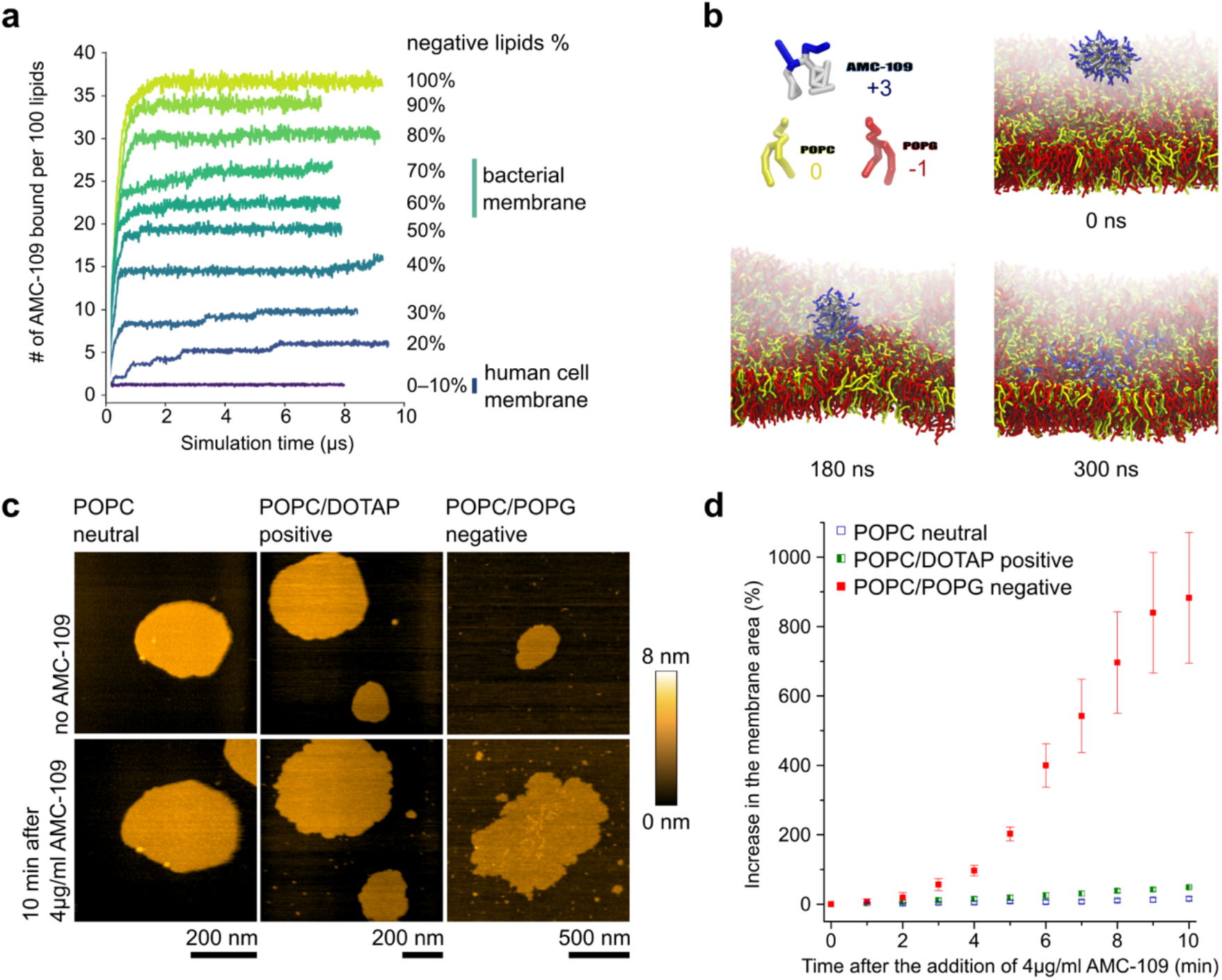
Interaction of AMC-109 aggregates with model membranes. **a** MD results on the number of AMC-109 molecules attached to the membrane per 100 lipids. An increase in a curve corresponds to the attachment of one or more aggregates into the membrane. The curves for 0 and 10% of negatively charged lipids both show no attachment of the AMC-109 aggregates. Levels of the negative charge corresponding to the membranes of human and bacterial cells are highlighted. **b** Interaction of a single AMC-109 aggregate with a symmetric POPC/POPG (40/60 mol%) membrane. The aggregate attaches to the membrane and gradually all the AMC-109 monomers dissolve in between the lipids. The simulation was run with multiple AMC-109 aggregates (Figure S8). Other AMC-109 aggregates and water are hidden for clarity. Time stamps refer to the time of the simulated MD trajectory. **c** Representative HS-AFM images of the synthetic lipid membranes before and 10 minutes after the addition of 4 μg/ml AMC-109. Experiments were performed on neutral POPC, positively charged POPC/DOTAP (40/60 mol%), and negatively charged POPC/POPG (40/60 mol%). Growth of supported membranes deposited on mica was monitored in time after the addition of the AMC-109. **d** Quantitative analysis of the increase in the surface area of synthetic lipid membranes in time after treatment with 4 μg/ml AMC-109. Data correspond to representative images shown in panel **c**. Negatively charged membranes prove to be significantly more affected than neutral and positively charged ones. Averages and standard errors of the mean measured for each lipid composition for 3 individual experiments on 6–7 membrane patches are shown.

The insertion of a single AMC-109 aggregate is depicted in Figure 4b. First, the aggregate makes direct contact with the negatively charged lipid headgroups in the membrane (red in Figure 4b), then it remodels the upper leaflet of the attacked membrane pushing the lipids from each other and inserting the AMC-109 molecules from the whole aggregate in between the phospholipids. Interestingly, the AMC-109 molecules from a single aggregate enter only the one leaflet of the membrane and do not translocate into the opposite one within the simulation time scales. Once inserted into the hydrophobic interior of the membrane, AMC-109 molecules move as monomers, and do not further cooperate towards any membrane disruption. The whole process of the aggregate adsorption and dissolution into the membrane takes ~300 ns in the simulation. The time needed for the saturation of the membrane with the AMC-109 molecules is ~2 μs for membranes with higher content of the negative charge (> 30% POPG, Figure 4a), and longer (> 5 μs) for membranes less attractive for AMC-109 (≤ 30% POPG).

We further test the selectivity of the aggregates towards negatively charged membranes by HS-AFM experiments (Figure 4c,d), where we observe whether expansion of synthetic lipid membranes occurs in the first 10 minutes after exposure to 4 μg/ml AMC-109. Neutral (POPC) and positively charged membranes (DOTAP/POPC, 60/40 mol%) increase their surface area by 16 ± 5% and 49 ± 3%, likely due to incorporation of few AMC-109 monomers from the solution. The negatively charged (POPG/POPC 60/40 mol%) grows by 880 ± 190%. The significantly higher effects to the negatively charged membranes can be explained by the additional incorporation of the micellar aggregates, as predicted by the MD simulations.

Taken together, the selectivity of AMC-109 to bacteria seems to arise from two main factors. First, the amphiphilic nature of AMC-109 drives formation of nanometer-sized aggregates hiding their hydrophobic parts and exposing their cationic groups. MD simulations agree on the shape and characteristic dimensions of such aggregates with HS-AFM images, which also demonstrate their existence at concentrations around the MIC for *S. aureus*. Second, self-assembly of AMC-109 creates an energetic barrier^37^ preventing interaction of AMC-109 with neutral membranes lowering the cytotoxicity of the compound for human cells^39,41^. Both MD simulations and HS-AFM experiments show that this barrier is lowered by the bacterial membrane surface charge, which also facilitates attraction of the aggregates. This leads to their loading into the bacterial membrane resulting in severe damage of the membrane lateral organization, which we suggest to be the cornerstone of the antimicrobial activity of AMC-109. Potentially, this could even have implications for cancer treatment as multiple studies already investigated the possibility of therapeutic application of cationic peptides^39–41^.

### AMC-109 does not form membrane pores

Our results suggest that AMC-109 incorporates into the bacterial membrane and changes the lateral lipid organization dissolving the lateral domains, which changes both dynamics of the lipids and the membrane material properties. Neither simulations, nor experiments reveal any porous defects in the membrane. To confirm that we did not overlook any small membrane perturbations, we performed cobalt-calcein leakage assays and compared the leakages induced by AMC-109 and by the well-known pore forming peptide melittin (Figure 5). AMC-109 induces no leakage of calcein in POPG/POPC/POPE (60/30/10 mol%) liposomes up to very high concentrations of 6 AMC-109 molecules per single lipid (600 μg/ml AMC-109 added to 100 μg/ml liposomes). Only a these very high concentrations of AMC-109, we observe slow but continuous leakage without any saturation behavior (Figure 5b), which corresponds to the overall disruption of liposomes^42,43^. In contrast, pore forming melittin induces pores in the liposomes yielding a gradual leakage corresponding to the creation of individual pores (Figure 5c)^42,44,45^.

**Fig. 5.**
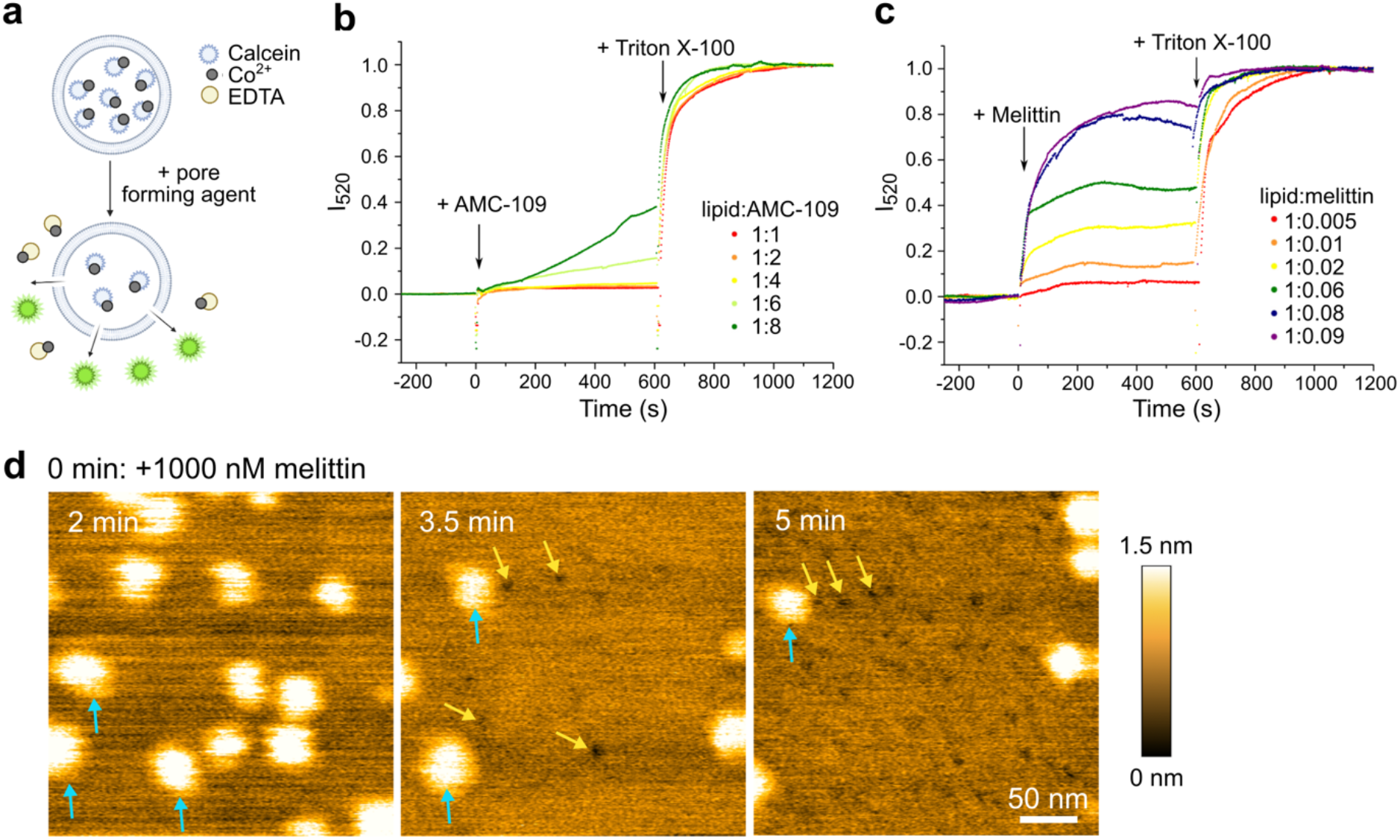
Pore-forming activity of AMC-109 and melittin. **a** Cobalt-calcein leakage assays principle. Non-fluorescent cobalt-calcein complex is trapped inside of the membrane liposomes. The addition of a perforating agent leads to their leakage out of the liposomes, where cobalt is extracted from the complex by EDTA resulting in an increased calcein fluorescence intensity. **b,c** Calcein leakage induced by AMC-109 and melittin in POPG/POPC/POPE (60/30/10 mol%) liposomes. Arrows indicate the addition times of the antibiotic agent and detergent Triton X-100 to fully rupture the liposomes and obtain maximum fluorescence signal. 100 μg/ml liposomes were used for the experiments. **b** AMC-109 does not induce any leakage up to a ratio of 6 AMC-109 molecules per single lipid. **c** Melittin displays gradual leakage corresponding to its pore forming activity. **d** HS-AFM images of *S. aureus* lipid membranes after the addition of 1000 nM melittin. Blue arrows point at the lateral domains, visible as bright parts of the membrane. Yellow arrows point at the formed pores. Initially a lot of lateral domains are visible (left), pores start appearing after a few minutes of melittin exposure (middle), and gradually more pores are being formed (right).

The striking differences in membrane pore forming activity of melittin and the lack of pore formation after AMC-109 treatment are visualized by HS-AFM (Figure 5d). While AMC-109 largely affects lateral domain organization (Figure 2) but does not induce detectable membrane pore formation (Figure 1–2, and 5b), adding melittin leads to the formation of individual small pores (yellow arrows in Figure 5d, Figure S9). Such pores, with diameters below 10 nm, are distributed over the whole membrane surface (Figure 5d, Figure S9). Melittin also slows down the lateral movement of the lipid domains, and some of the domains disappear (Figure 5d). In summary, these observations support our conclusion that AMC-109 does not perforate the *S. aureus* membrane in a manner comparable to pore-forming peptides but instead acts by affecting its lateral organization.

### AMC-109 acts like a bacteria-selective disinfectant

The mechanism of the activity of AMC-109 hence does not resemble that of pore-forming peptides, or of other membrane active antibiotics, which perforate the membranes via various types of pores or carpet-like membrane coverage and disruption^3,46,47^. Incorporation of AMC-109 molecules in between the lipids of a single membrane leaflet and effects on lateral organization in fact resemble the activity of small disinfectant molecules like ethanol, benzalkonium chloride (BAK) or benzyl alcohol ^48–58^.

Ethanol (Figure 6a) was previously shown to thin model lipid membranes, lower the order of the lipid tails, and dissolve lateral cholesterol-rich domains in membranes^49–52^. BAKs are molecules containing a quaternary nitrogen associated with hydrophobic headgroup and a single hydrophobic acyl chain of a varying length (Figure 6). These surface-active molecules are believed to insert in between the lipids inside of the membrane, disturb the lateral organization, and — in high concentrations — form mixed lipid/BAK micelles tearing out parts of the cytoplasmic membrane^53–56^.

**Fig. 6.**
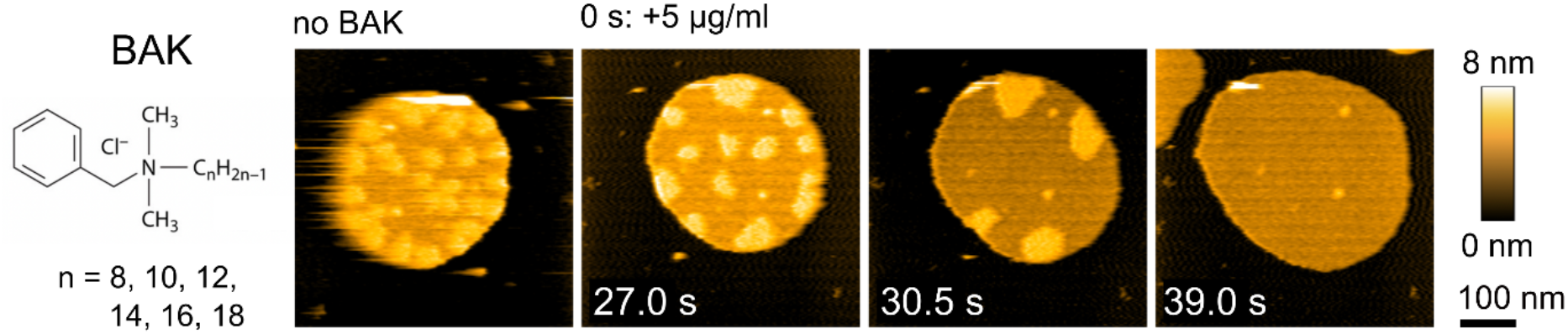
Effects of the disinfectant benzalkonium chloride (BAK) on *S. aureus* lipid membranes. Chemical structure of BAK and HS-AFM images of the membrane before and after the addition of 5 μg/ml BAK.

We performed HS-AFM imaging of the *S. aureus* lipid membranes exposed to BAK (Figure 6) and ethanol (Figure S10). For BAK (Figure 6), we observe activity strikingly similar to AMC-109. Domains are clustered, which is followed by their dissolution and expansion of the membranes. From the molecular point of view, both AMC-109 and BAK thus seem to similarly incorporate in between the individual lipids in the *S. aureus* lipid membranes and disturb its lateral organization clustering and dissolving the lateral domains. In the case of ethanol we observe slowing down of the lateral movement of the domains and their fast dissolution. The effects of ethanol are very fast (Figure S10), which points to a low kinetic barrier for incorporation of the small ethanol molecule into the lipid membrane. We hypothesize that the bigger BAK molecules have to orient themselves prior to insertion into the membrane. Similarly, AMC-109 have to dissolve their aggregates before entering the membrane. For both AMC-109 and BAK, this represents a kinetic barrier making them acting slower than ethanol.

BAK has been used widely for more than 70 years as a disinfectant and an antimicrobial in agriculture and industrial production including for instance antimicrobial soaps^59^. It’s widespread use led to partial reduced susceptibility in bacteria but full resistance has not been observed. Disinfectants are indeed powerful molecules, however, they cannot be used as antibiotics in human medicine for their toxicity to eukaryotic cells. AMC-109 with its high selectivity for bacteria is not observed to be toxic, and the similarities with the BAK and ethanol mode of actions suggest that for AMC-109 it might take a while before resistance is generated in bacteria.

## Conclusions

In conclusion, AMC-109 uniquely combines multiple mechanisms in the fight against bacteria. AMC-109 monomers first form stable aggregates, which selectively interact with the negatively charged surface of bacteria. AMC-109 in its monomeric form is highly hydrophobic so it tends to interact with all membranes including those of mammalian cells. The aggregation, in which the hydrophobic part of the AMC-109 molecule is hidden inside the aggregate and only the positively charged residues are exposed to water, is thus a crucial step responsible for the drug’s high selectivity for bacteria and the absence of toxicity for human cells. The aggregates then attach to the bacterial cytoplasmic membrane and gradually dissolve all the AMC-109 individual molecules into the outer membrane leaflet. Once inside the membrane, AMC-109 affects the membrane lateral organization and lipid order, dissolving membrane domains. Functional membrane microdomains (FMMs) are important in *S. aureus* cells for protein sorting, signaling, or cell wall synthesis, as well as for the resistance of MRSA for penicillin, methicillin and other β-lactam antibiotics. Their dissolution takes all these functionalities away, leads to overall stiffening of the membrane, and possibly makes the MRSA colony again susceptible to β-lactams. The fact that AMC-109 targets negatively charged lipids, a basic feature of bacterial membranes, together with its similarities with the activity of the disinfectant BAK makes this antimicrobial agent particularly promising in not generating resistance in bacteria for a longer time. The high selectivity of AMC-109 for bacteria over human cellular membranes ^11^, provided by its aggregating properties, at the same time supports safer usage in medical treatment. We performed study on membranes from *S. aureus* lipid extracts, however the findings on the AMC-109 activity could be applicable to a wide range of Gram-positive bacteria, as well as cancer cells, which also possess a net negative surface charge. By deciphering the molecular mode of action of AMC-109 we identified lateral membrane organization as a viable drug target, which can be used as a target in further pharmaceutical development. Dissolving membrane domains makes AMC-109 a highly selective disinfectant that is active against *S. aureus* bacteria including multi-resistant strains and possibly makes them again susceptible to treatment by β-lactam antibiotics like penicillin or methicillin.

## Acknowledgments

WHR thanks the EU for support through the INFRAIA infrastructure grant MOSBRI. AM thanks the Nederlandse organisatie voor Wetenschappelijk Onderzoek (NWO) for support through a Physics/f grant (no. 680-91-007) and an XS grant (no. OCENW.XS22.1.083).

## Author contributions

AM, SM, JM and WHR have made a substantial contribution to the concept and design of the work; AM, SM, JM, NAWdK, MG, and Jv/dE took part in the data collection, data analysis and interpretation; AM drafted the article; AM, SM, JM, NAWdK, WS, JSMS, AJMD, SJM, and WHR have critically revised the article; All the authors have approved the final version to be published.

## Methods

### Materials

AMC-109 (previously LTX-109^11^; Figure 1a) was obtained in the form of dry powder from Amicoat AS, Norway. The powder was dissolved in miliQ water to stock solutions of 1–25 mg/ml, stored at room temperature shielded from the access of air and light, and used for the maximum of one month.

*Staphylococcus aureus* cells, strain RN4220, NCTC8325-4 derivative, restriction deficient and cured of prophages^60^, was a generous gift from prof. J. M van Dijl from University Medical Centrum Groningen. The cells were stored in −80°C.

Phosphate buffered saline (PBS buffer, pH 7.4) in the form of tablets for dissolution in miliQ water was purchased at Sigma-Aldrich. Synthetic lipids 1-palmitoyl-2-oleoyl-sn-glycero-3-phospho-(1’-rac-glycerol) (POPG), 1-palmitoyl-2-oleoyl-glycero-3-phosphocholine (POPC), and 1-palmitoyl-2-oleoyl-sn-glycero-3-phosphoethanolamine (POPE), di-oleoyl-phosphatidylglycerol (DOPG), di-oleoyl-biphosphatidylglycerol cardiolipin (DODOCL) and/or mono- and di-galactosyldiacylglycerol (MGDG and DGDG) were purchased from Avanti Polar Lipids as 25 mg/ml solutions in chloroform.

Pore forming peptide melittin was purchased from GenScript Biotech (Netherlands) B.V. in the form of dry powder. The powder was dissolved in DMSO to stock concentration of 25 mg/ml and stored in −20°C. The stock solution was diluted in miliQ water to 1 mM solution and used for one day. Final DMSO concentration during the AFM and leakage experiments was < 0.010 vol% and < 0.015 vol%, respectively. Test experiments showed no effect of DMSO on the membranes at these concentrations.

Benzalkonium chloride, a mixture of molecules with acyl chain length varying between 8 to 18 carbons, was purchased in the form of semisolid gel with purity ≥ 95% from Sigma-Aldrich. The semisolid gel was dissolved in miliQ water to the stock concentration of 1 g/ml. Ethanol with purity ≥ 97% was purchased from Boom B.V.

For the lipidomic studies: ammonium formate, 1-butanol, alpha-naphtol, molybdenum blue and ninhydrin were purchased from Sigma-Aldrich. Acetonitrile was purchased from Biosolve (Netherlands) B.V. For cobalt-calcein leakage assays: calcein, ethypenediaminetetraacetic acid (EDTA), H_2_KPO_4_, HK_2_PO_4_, and CoCl_2_ hexahydrate with purity >98%, and the filtration gel Sephadex G75 were purchased from Sigma-Aldrich.

### *S. aureus* cells cultivation

*S. aureus* cells were cultivated in a growth medium in 37°C, while shaking at 200–250 rpm, until the saturated growth phase. The cells were centrifuged at 2095 × g (centrifuge Allegra X-15R by Beckman, swing-out rotor SX4750A) at 4°C for 10 min and the growth medium was exchanged for PBS buffer. The centrifugation step was repeated again and the excess PBS buffer was taken out. Wet cells were stored at −80°C until lipid isolation.

### Lipid isolation from *S. aureus* wet cells

The lipids were extracted following the modified Bligh & Dyer extraction protocol^17^. In short, the cells were first washed in miliQ water and the pellets were transferred into tubes and weighted. The cells were dissolved in water/chloroform/methanol in ratios 0.8:1:2, where the water content was 0.8 ml per 1 g of the cells, and left stirring in 4°C overnight. The mixtures were then centrifuged at 900 × g at 4°C for 15 min. The supernatant was resuspended in a 1:1 chloroform/water mixture and left to phase separate at room temperature for 2 days. The bottom layer was collected and dried in a rotary evaporator. Finally, the dry lipid film was weighed and dissolved in chloroform to the stock concentration of 10 mg/ml and stored at −20°C.

### Lipidomic analysis of the *S. aureus* lipid extracts

Lipid extracts were analyzed by UHPLC-MS for the determination of the presence of various lipid species and their saturation state as described in detail elsewhere ^61,62^. In brief, 5 μl of a lipid extract (10 mg/ml in methanol) was injected on a Waters Acquity CSH C18 column. Lipids were separated with a gradient elution starting at 55% mobile phase A (5 mM ammonium formate in water:acetonitrile, 40:60) and 45% mobile phase B (5 mM ammonium formate in water:acetonitrile:1-butanol, 0.5:10:90) and by increasing B to a maximum value of 90% over 19.5 minutes. The resulting mass spectra were analyzed using the Thermo Xcalibur Qual Browser (Thermo Fisher Scientific).

Relative quantification of glycerophospholipids and glycolipids was performed by spotting 1, 5 and 10 μl of the lipid extract 2 cm from the bottom on aluminum-backed silica gel 60 TLC plates (cat#: 1.05553.0001; Merck) together with DOPG, DODOCL and/or MGDG and DGDG reference compounds. Lipids were separated using a mobile phase consisting of chloroform:methanol:glacial acetic acid (65:25:8) until the solvent front reached a point approximately 1 cm from the top of the TLC plate. The TLC plates were first dried at RT and then stained by momentarily horizontally dipping them in a shallow pool of staining solution: (a) molybdenum blue staining solution for the visualization of phosphate-containing lipids, (b) 0.5% (w/v) α-naphtol in ethanol or 1-butanol with H_2_SO_4_ (9:1, v/v) (adapted from ^63^ to visualize glycolipids or (c) 0.1% (w/v) ninhydrin in water-saturated 1-butanol^64^ to visualize lipids containing primary or secondary amines. The TLC plates were then left at room temperature to develop and heated with a hairdryer if necessary.

For relative quantification, the TLC plates were digitized using a PowerLook 1120 scanner (Amersham Biosciences) in combination with the VueScan software (v9.094, Hamrick Software). Images were analyzed using the Fiji^65^ “gel analyzer” tool (build: ImageJ V1.53c) to produce chromatograms and calculate integrated peak values of which relative values were calculated.

### Atomic force microscopy experiments

Preparation of liposomes: *S. aureus* lipid extracts in chloroform were pipetted into the glass vial. The chloroform was evaporated using an argon stream and let dry completely for >1 hour in a vacuum. PBS buffer was added to the lipid film in the concentration of 0.3 mg/ml. To help the resuspension, the mixture was vigorously shaken for 1 minute followed by five cycles of freezing by liquid nitrogen and thawing in warm water. The resuspended liposomes were then extruded 21 times through 0.1 μm pores polycarbonate membranes (Avanti Polar Lipids). The liposomes were stored in 4°C and used for the maximum of one week.

Sample cell for AFM was prepared by gluing a small piece of mica on microscope glass slide with a transparent epoxy glue. A glass ring was attached around it by two component biocompatible glue (Bruker Nano GmbH). The liposomes were diluted to 0.02–0.04 mg/ml in PBS buffer. 10 μl drop of the diluted liposomes was deposited on freshly cleaved mica surface and let to sediment for >10 min. In some cases, the PBS buffer was washed out 5 times and replaced with a new 10 μl drop of PBS buffer to press the liposomes on the mica surface and support the rupture of the liposomes and formation of isolated supported membrane patches. PBS buffer was then added to the total volume of 1 ml.

Supported membranes made of *S. aureus* lipid extracts are denoted as *S. aureus* lipid membranes in the text.

AFM imaging and nano-indentation of the supported *S. aureus* lipid membranes immersed in PBS buffer was performed with an JPK Nano Wizard Ultra Speed AFM. The experiments were performed at room temperature (22°C) using qp-BioAC cantilevers (NanoAndMore GmbH) with a nominal spring constant 0.06 ± 0.03 N/m and a silicon nitride tip with a typical tip radius of curvature smaller than 10 nm. The imaging force was ~80–100 pN in all cases.

During the nano-indentation experiment, the AFM tip approaches the supported membrane in a constant velocity and indents the membrane until reaching the underlaying support (Figure S6a). First, the membrane was imaged, then the hard mica surface was indented with a force up to 2 nN with an indentation velocity of 300 nms^−1^ to check for any tip contamination and get a hard surface reference, which gives us the bending behavior of the cantilever itself. Immediately after the check of the tip contamination the membrane was indented with a force up to 1 nN with an indentation velocity of 300 nms^−1^. The dependence between the force applied on the membrane *F* and the vertical tip position *z—*the force-indentation curve—is recorded. If multiple spots on membrane were indented, clean surface curve was measured before each new membrane indentation. After indentations, the same spot was imaged again. For AMC-109 concentrations > 2 μg/ml, in which the membrane expands over the substrate, nano-indentation experiments were not possible due to adhesion forces in between the AFM tip and the surface. This was probably due to AMC-109 molecules covering both the expanded membranes and the surrounding mica surface. We hence focused our investigation on the mechanical properties of *S. aureus* lipid membranes exposed to AMC-109 concentrations < 2 μg/ml. At 1 μg/ml in part of the membranes the lateral domains are completely dissolved, whereas in some membrane patches they are still visible. There is also a part of the membranes where we couldn’t make a clear distinction between these two cases. Moreover, it is not possible to reliably distinguish from the force-indentations curves if we are indenting a region on top of the domain or out of it. Mechanical properties shown in Figure 3 were, hence, calculated from all data, gathered both on top and out of the lateral domains.

Both the AFM images and the force curves were processed using JPK Data Processing Software. AFM force-indentation curves analysis: In brief, the nano-indentation experiment yield the deflection of the cantilever as a function of the Z-piezo displacement. This dependency was converted to the applied force versus the vertical distance between the tip and the sample surface by subtracting the deflection of the cantilever itself^66^, which was determined from the force-distance curves recorded on a hard surface. The penetration point, when the tip perforated the membrane was determined as a point where we observe an apparent drop in the applied force. The region between the first tip–membrane interaction and the full membrane penetration was fitted with a modified Hertz model using Eq. S1 according to a modified Hertz model of a thin supported layer with a 1 nm water layer bellow^50^. Two parameters were derived from the fit: vertical distance of the first tip– membrane interaction related to the membrane height and Young’s modulus of the membrane describing its tensile stiffness. The analysis of the mechanical properties of the *S. aureus* lipid membranes in the presence of 0, 0.5 and 1 μg/ml AMC-109 was done independently on 150, 119, and 144 individual force-indentation curves, respectively. Resulting Young’s modulus values (Figure S5) are reported as median ± standard error of the mean.

### High-speed atomic force microscopy

Liposomes made from *S. aureus* lipids were prepared for the HS-AFM experiments the same way as for AFM. Synthetic liposomes were prepared in concentrations 1 mg/ml in PBS. In cases where attachment of membranes to mica was weaker, we exchanged the last extrusion step in the liposomes preparation with gentle sonication for 30 s in a sonication bath prior to the liposomes deposition on mica. Expansion of synthetic lipid membranes was measured at POPC, POPC/DOTAP (40/60 mol%), and POPC/POPG (40/60 mol%). Negatively charged membranes made of only two components could not be reliably attached on mica, so we opted for a more complex composition fitting our lipidomic analysis of the *S. aureus* lipid extracts.

HS-AFM experiments were done using RIBM (Japan) machine in amplitude modulation tapping mode in liquid^23,24,67,68^. Short cantilevers USC-F1.2-k0.15 (NanoWorld, Switzerland) with a spring constant of 0.15 N/m, resonance frequency around 0.6 MHz, and a quality factor of ~2, and ultra-small cantilevers BL-AC10FS-A2 (Olympus, Japan) with a spring constant of 0.1 N/m, and resonance frequency around 0.6 MHz in buffer were used. The cantilever free amplitude was set to 1 nm, and the set-point amplitude for the cantilever oscillation was set around 0.8 nm. Images were taken at 300 ms to 6 s per frame depending on the size of the image. A mica surface of diameter 1.5 mm glued on top of a 5 mm high glass rod was used as the AFM sample stage. The glass rod was then attached to the scanner Z-piezo using a small amount of nail polish. For the activity of the antibiotic agents (AMC-109, melittin, ethanol, BAK) on synthetic and *S. aureus* lipid membranes, a 3 μl drop of 0.03–0.5 μg/ml liposomes in PBS buffer was deposited on the freshly cleaved mica and the buffer was washed out 5 times to press the liposomes on the mica surface and support the rupture of the liposomes and formation of isolated supported membrane patches (as in AFM experiments). The scanner head was then put upside down into a small liquid chamber containing the cantilever and filled with 40–80 μl of the recording solution (PBS buffer). First, we image untreated membranes and then add 5 μl of the buffer with a concentrated solution of the active agent to achieve the desired final concentration in the sample.

Size distribution of lateral lipid domains was measured at 269 individual domains across multiple experimental days. They showed diameter of 31 ± 6 nm. Membrane thickness was measured by comparison of the measured height at the parts outside of the lateral domains with the height measured at the adjacent mica surface. Similarly, height of the AMC-109 aggregates was measured as a difference between the height measured on top of the aggregate and on the surrounding mica. Expansion of the synthetic lipid membranes after the addition of 4 μg/ml AMC-109 was measured from 3–4 individual experiments per lipid composition, in which expansion of 7–14 membrane patches was analyzed. Height, thickness, membranes expansion, and domains size measurements from AFM and HS-AFM images are reported as mean ± standard error of the mean.

### Fluorescence leakage assays

In cobalt-calcein fluorescence leakage experiments, calcein is trapped inside of the membrane liposomes, where its fluorescence is quenched by formation of complexes with cobalt ions (Figure 6a). The addition of a perforating agent to the outside buffer leads to the leakage of cobalt-calcein complexes out of the liposomes, where cobalt is extracted from the complex by ethylenediaminetetraacetic acid (EDTA) resulting in an increased calcein fluorescence intensity.

50 mM and 35 mM phosphate buffers, pH 7.0, were prepared as mixtures of H_2_KPO_4_ and HK_2_PO_4_. 35 mM phosphate buffer was mixed with 10 mM EDTA and pH was adjusted to pH 7.0. POPG/POPC/POPE lipids in ratios 60:30:10 mol% were pipetted from the stock solutions in chloroform into the glass vial. Chloroform was evaporated using argon stream and let to completely dry out for >1 hour in vacuum. The lipid film was rehydrated using a mixture of 1 mM CoCl_2_ and 0.8 mM calcein dissolved in 50 mM phosphate buffer, pH 7.0, into the lipid concentration of 100 mg/ml. Resuspension was promoted by five cycles of freezing in liquid nitrogen and thawing in warm water. Formed liposomes were extruded 21 times through 0.4 μm pores polycarbonate membranes (Avanti Polar Lipids). The extruded liposomes were stored in 4°C overnight.

On the day of the experiment, the extruded liposomes were washed from the external buffer containing free CoCl_2_ and calcein. The original buffer was replaced with the 35 mM phosphate buffer with 10 mM EDTA, pH 7.0 using gravity separation column with the internal volume of 12 ml packed with Sephadex G-75. The column was first preequilibrated with the 35 mM phosphate buffer with 10 ml EDTA, pH 7.0, and then used for separation of the extruded liposomes from the original external buffer and the free calcein and CoCl_2_. All fractions containing slightly fluorescent liposomes were collected into a single tube and ultracentrifugated at 444 000 × g for 25 min at 4°C. The buffer on top of the pellet of liposomes was removed, and the pellet was resuspended in fresh 35 mM phosphate buffer with 10 mM EDTA, pH 7.0 to the stock concentration of 100 mg/ml.

The fluorescence measurements were performed with Fluorometer QuantaMaster 40 (PhotoMed GmbH). The liposomes were diluted to 0.1 mg/ml for the experiments. The calcein was excited at λ_ex_ = 495 nm and the emission at λ_em_ = 520 nm was detected in time with both the excitation and emission slits opened to the range of wavelengths ± 5 nm. Background fluorescence was measured for 5 min, then membrane-active agent (AMC-109 or melittin) was added and the increase in the fluorescence intensity from the calcein leaking out of the liposomes was detected for 10 min. Finally, detergent Triton X-100 in the final concentration of 0.25 vol% was added to rupture all the liposomes and release all of the calcein. The maximum intensity of the fluorescence signal was detected for 10 min. The time traces of the fluorescence intensity were normalized to 0 at the background intensity before the addition of the membrane-active agent and to 1 at the maximum intensity after the addition of Triton X-100.

### Molecular dynamic simulations

The initial configurations of the simulated lipid bilayers were generated using the tool Insane^69^ to yield lateral dimensions of 30×30 nm from 3042 lipids in total with the bilayer repeat distance of 30 nm. Using the GROMACS tool *gmx solvate*^70^, 2489 AMC-109 molecules were distributed in a water solution with a minimum distance of 1 nm between each of them. The resulting molar ratio of AMC-109:water molecules was 1:250 (approximately 4 water molecules are represented by 1 Martini bead). The total charge of the system is neutralized by Chloride counterions with an additional 100 mM NaCl concentration in the solution.

All systems were equilibrated for at least 100 ns prior to production simulations. A standard simulation setup for the Martini model was used as described previously^35^. In brief, the simulation temperature was coupled to a v-rescale thermostat^71^ at room temperature of 310 K, separately for lipids, AMC-109 and solvent. A Parinello-Rahman barostat^72^ was used for pressure coupling at 1 bar with a coupling constant of 24 ps independently for the membrane plane and its normal. Standard time step of 20 fs was used for all simulations and the trajectory was recorded every 1 ns. For production, we have simulated all systems for at least 7 μs to ensure convergence.

The topology for AMC-109 was first generated based on the corresponding peptide template, using the protocol outlined in ref. ^35^ and the resulting peptide topology was subsequently modified to include the modification in the Tryptophan residue and the changes to the C-terminus as outlined in ref. ^11^. The topologies for lipids, ions and solvent were taken directly from the Martini 3 collection^35^. Simulations were run using GROMACS simulation package ver. 2019.3 in a mixed precision compilation without GPU support^73^. The profiles of the number of contacts between AMC-109 and lipids was calculated using GROMACS tool *gmx mindist* with a cutoff radius of 0.6 nm between the groups of molecules. Scripts used to prepare, perform and analyze the simulations, also including molecular topologies, simulation setup files and initial configurations are available in an open public repository https://github.com/jmelcr/peptidomimetics.

