## Supplementary Information for "Lateral membrane organization as target of an antimicrobial peptidomimetic compound"

**List of Supplementary Approaches:**

Nano-indentations of *S. aureus* lipid membranes: Dissolution of lateral lipid domains leads to changes in membrane elasticity

**List of Supplementary Figures:**

Figure S1: Lipidomic analysis of *S. aureus* lipid extracts

Figure S2: Expansion of the *S. aureus* lipid membranes after the treatment with 2 and 3 µg/ml AMC-109

Figure S3: 1x1 µm view of the changes in the *S. aureus* lipid membranes induced by the treatment with AMC-109

Figure S4: Diameter of the lateral lipid domains in untreated *S. aureus* lipid membranes

Figure S5: Nano-indentations of the *S. aureus* lipid membranes treated by AMC-109 in concentrations < MIC to evaluate membrane mechanical properties

Figure S6: Formation of a continuous carpet-like layer of AMC-109 on mica at HS-AFM setup.

Figure S7: Carpet-like layer of AMC-109 on mica surface.

Figure S8: Snapshot of the simulation box showing interaction of AMC-109 with the phospholipid membrane.

Figure S9: Melittin induced pores in *S. aureus* lipid membrane visualized by HS-AFM.

Figure S10: Effects of ethanol on *S. aureus* lipid membranes observed by HS-AFM.

### Supplementary Approaches

#### Nano-indentations of *S. aureus* lipid membranes: Dissolution of lateral lipid domains leads to changes in membrane elasticity

We performed nano-indentation experiments[1–4] to evaluate the influence of the domains dissolution on the mechanical properties of the membrane (Figure S5). The nano-indentation data were fitted with a modified Hertz model, which encounters for a solid support below the membrane and ~1 nm water layer ($d_{WL}$) between the membrane and the support[2]:

$$F = \frac{16Y}{9} R^{\frac{1}{2}} (z - z_0)^{\frac{3}{2}} [1 + 0.884\chi + 0.781\chi^2 + 0.386\chi^3 + 0.0048\chi^4] \quad \text{(S1)}$$

where *R* is the radius of the AFM tip (typically 10 nm in our experiment) and $\chi = (R \times (z - z_0))^{1/2} / (z_0 - d_{WL})$. Young's modulus of the membrane *Y* and the vertical distance of the first tip–membrane interaction $z_0$ are the fitting parameters.

We focused on changes induced by AMC-109 at concentrations when the domains are accumulated or dissolved but the membrane is not yet expanded (0.5–1 µg/ml). The untreated *S. aureus* lipid membranes feature a double peak distribution of its thickness (Figure S5c) indicating a difference in height of the domains and the surrounding membrane. Upon the accumulation of the lateral lipid domains at 0.5 µg/ml AMC-109 the height differences between the two are less distinct as the thickness distribution appears to be broad but unimodal. Upon increasing the AMC-109 concentration to 1 µg/ml, we are at the edge concentration, when in some of the membrane patches the domains are accumulated, and in others the domains are already dissolved. These membranes display an increase in its Young's modulus from ~7 MPa for untreated membranes to ~17 MPa in the presence of 1 µg/ml AMC-109 (inset Figure S5b and S5f). Hence, dissolution of the lateral lipid domains and mixing of their content (likely cardiolipins and glycolipids[5,6]) with the surrounding membrane leads to overall stiffening of the membrane. This observation is in accordance with literature[7] which shows that cardiolipin in small concentration increases lipid lateral packing and also compressibility and mechanical moduli of membranes.

### Supplementary Figures

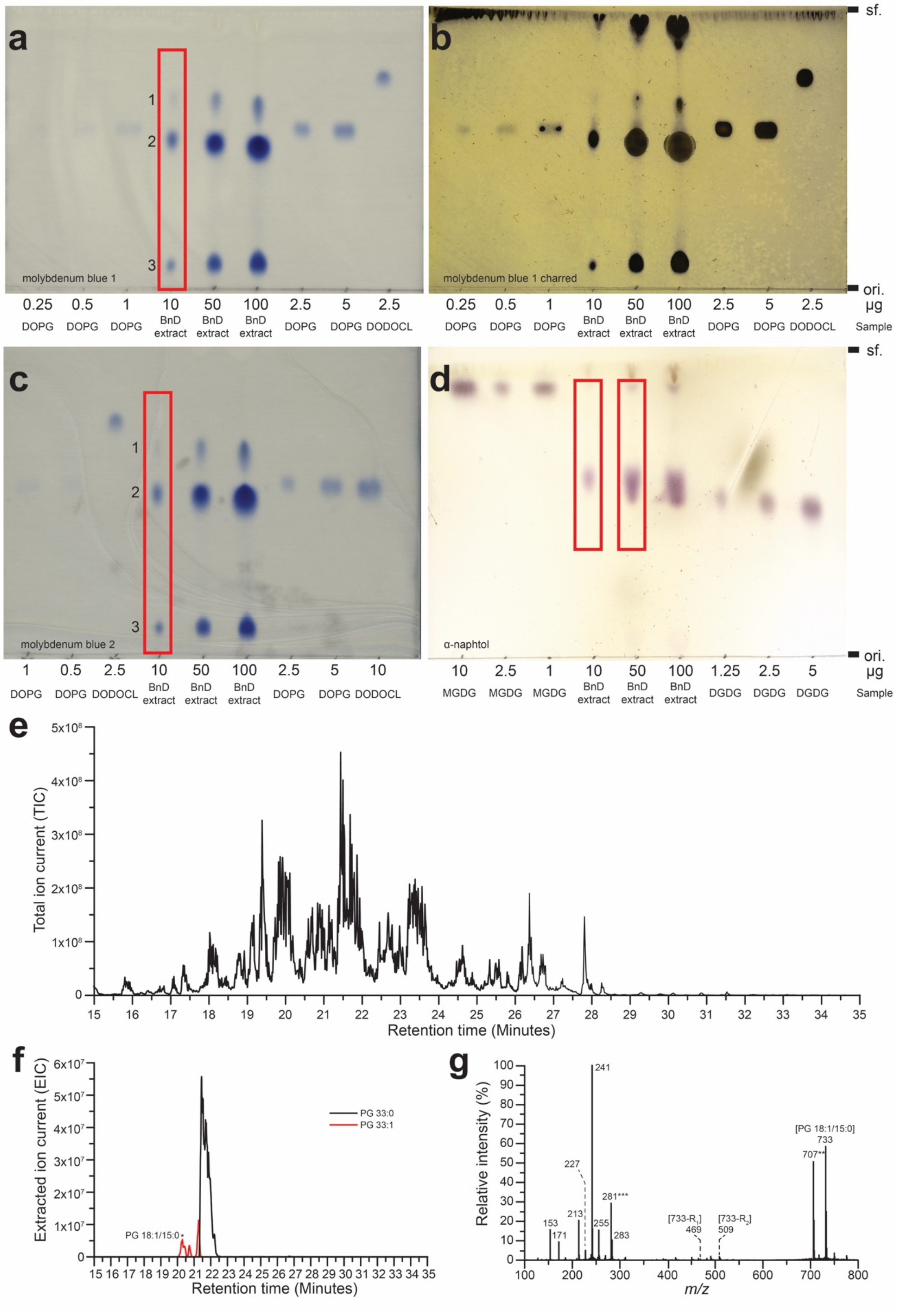

**Fig. S1**: TLC analysis plates (**a–d**) and Mass Spectra (**e–g**) of the *S. aureus* lipid extract. **a** and **c** Molybdenum blue stains indicate the presence of phosphate groups as blue spots. **b** Upon strong heating of a molybdenum blue stained plate unsaturated compounds char and show up as black spots while the blue spots of saturated phospholipids blends away in the background. Even a small amount of the unsaturated acyl tails results in a dark spot. Hence, no quantitative information about the content of the unsaturated phospholipids can be made. **d** α-naphtol stain indicates the presence of carbohydrate groups as purple spots. Red rectangles indicate the area on the TLC plate analysed by densitometry (**a**, **c** and **d**). **e** Total ion current (TIC) spectrum of the *S. aureus* lipid extract. (**f**) Extracted ion count for PG 33:0 from the spectra shown in (**e**). The most abundant PG phospholipid (black), and the unsaturated species PG 33:1 (red) are shown. **g** Fragmentation spectrum of PG 33:1 (18:1/15:0) (taken at time annotated by * in panel **f**) is shown with the parent ion (mass 733) and two second order fragmentation products (mass 469 and 503) which were used to assign its regioisomeric identity are annotated in brackets. ** Annotates the parent ion of a co-eluting phospholipid, PG 31:0. *** Indicates an unsaturated fatty acid. Unsaturated PG phospholipids were responsible for ~10% of the total PG phospholipid signal.

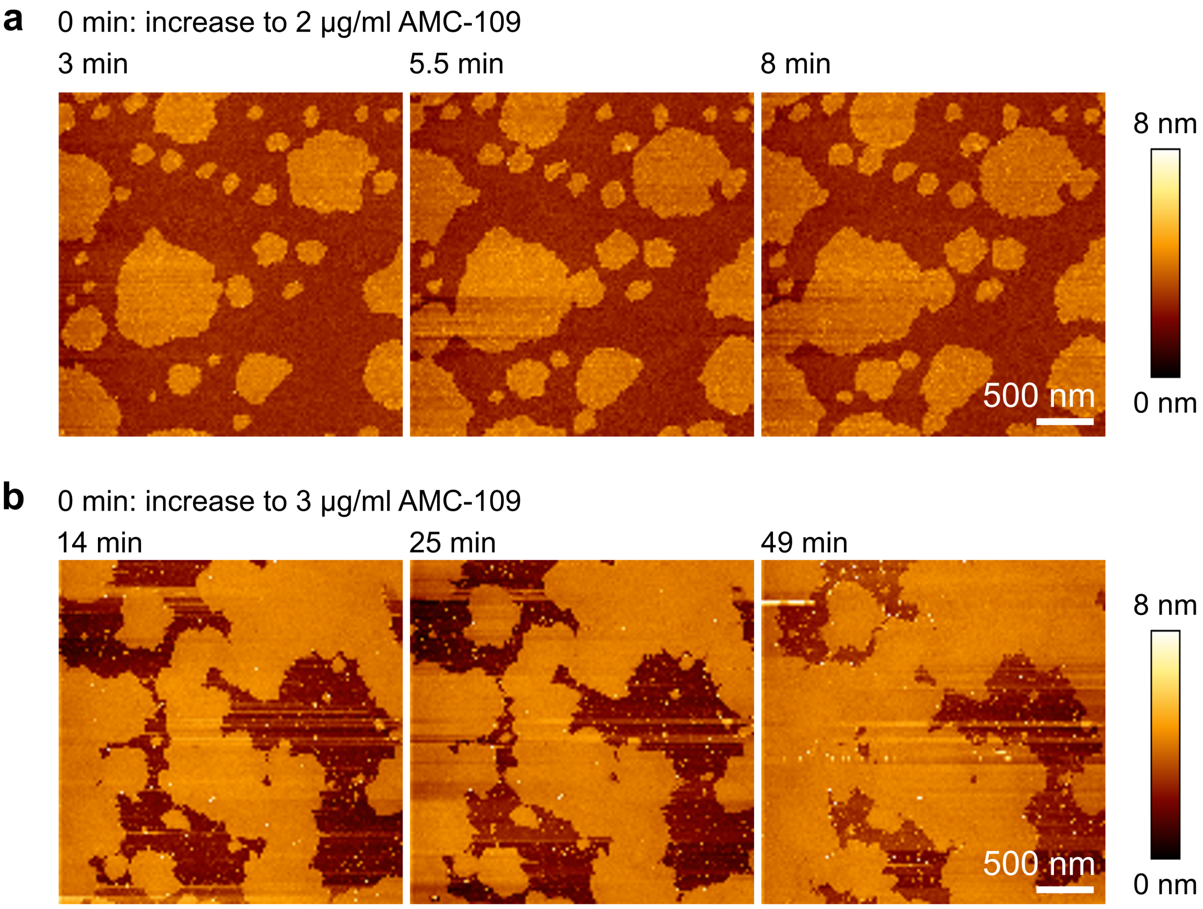


**Fig. S2**: Expansion of the *S. aureus* lipid membranes in time after the treatment with 2 (**a**) and 3 (**b**) µg/ml AMC-109. **a** 8 min after the addition of 2 µg/ml AMC-109 the system reaches equilibrium and the membrane stretching stops. In all measurements (N = 10) stretching stopped in <10 min. **b** At the concentration of 3 µg/ml, expansion is a continuous process and we see small changes even 49 minutes after the AMC-109 addition.

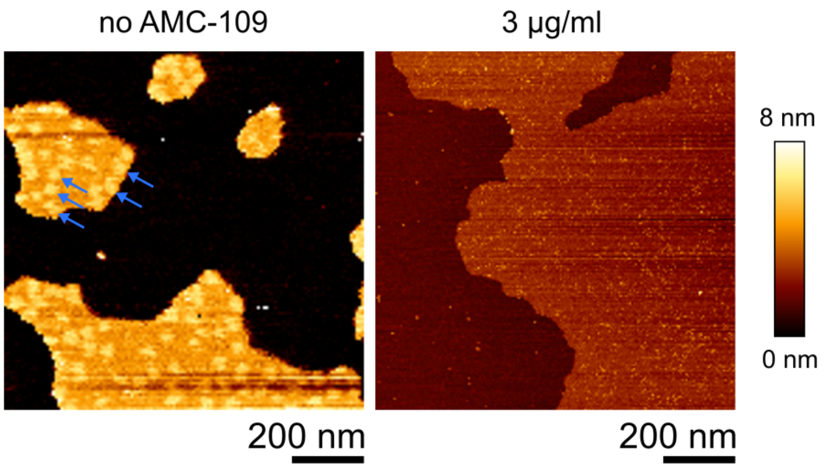


**Fig. S3**: 1x1 μm view of the changes in the *S. aureus* lipid membranes induced by the treatment with AMC-109. Blue arrows show lateral lipid domains in the untreated membrane. In the presence of 3 μg/ml AMC-109, i.e. above the minimal inhibitory concentration, the domains are lost, the membrane is expanded and thinned. The lighter color of the background indicates coverage of the mica surface with a layer of AMC-109 molecules.

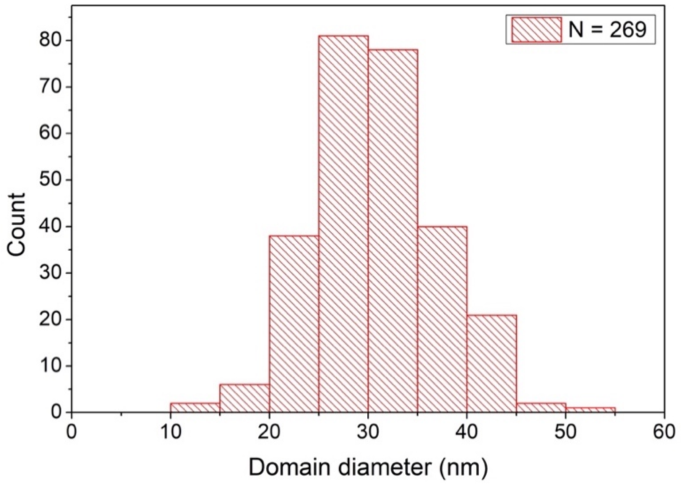


**Fig. S4**: Diameter of the lateral lipid domains in untreated *S. aureus* lipid membranes. Measured from HS-AFM images across four different experimental days. Average domain diameter is 30.9 ± 0.4 nm (N = 269).

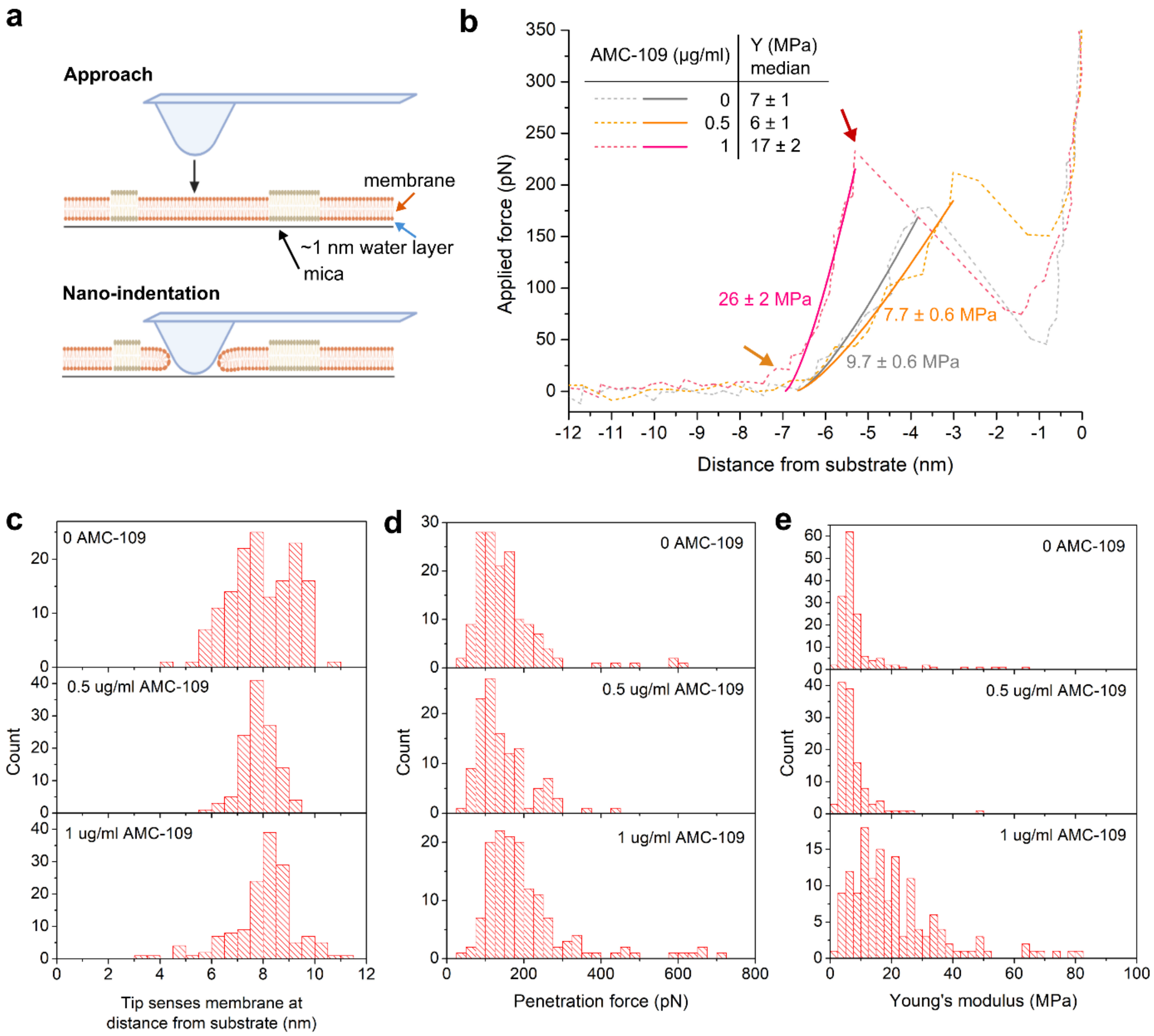


**f**

| AMC-109 | N total | Membrane height at tip interaction | Penetration force | Young's modulus |
|---|---|---|---|---|
| 0 | 150 | 8 ± 1 | 130 ± 10 pN | 7 ± 1 MPa |
| 0.5 µg/ml | 119 | 8 ± 1 | 120 ± 10 pN | 6 ± 1 MPa |
| 1 µg/ml | 144 | 8 ± 1 | 170 ± 20 pN | 17 ± 2 MPa |

**Fig. S5**: Mechanical characterization of *S. aureus* lipid membranes upon the interaction with AMC-109. **a** Schematics of the nano-indentation experiment. AFM tip approaches the supported membrane (orange arrow), which lays on the solid mica substrate (black arrow) with ~1 nm layer of water[8,9] (blue arrow) in between the membrane and the support. **b** Representative force-indentation curves in the presence of 0 (grey), 0.5 (orange), and 1 µg/ml (pink) AMC-109. The orange arrow indicates the first contact between the AFM tip and the membrane. The red arrow shows the point, when the tip penetrates through the membrane. Solid curves are the fits of the region in between these two points using the modified Hertz model for supported thin layers Equation S1[2]. Inset table features the median values and standard errors of Young's moduli of *S. aureus* lipid membranes in the presence of 0, 0.5, and 1 µg/ml AMC-109 (N = 150, 119, and 144, respectively). **c** Distance from the substrate of the first interaction between the tip and the membrane. **d** Force needed to penetrate the membrane. **e** Young's modulus of the membranes. **f** Median values and standard errors of the respective parameters.

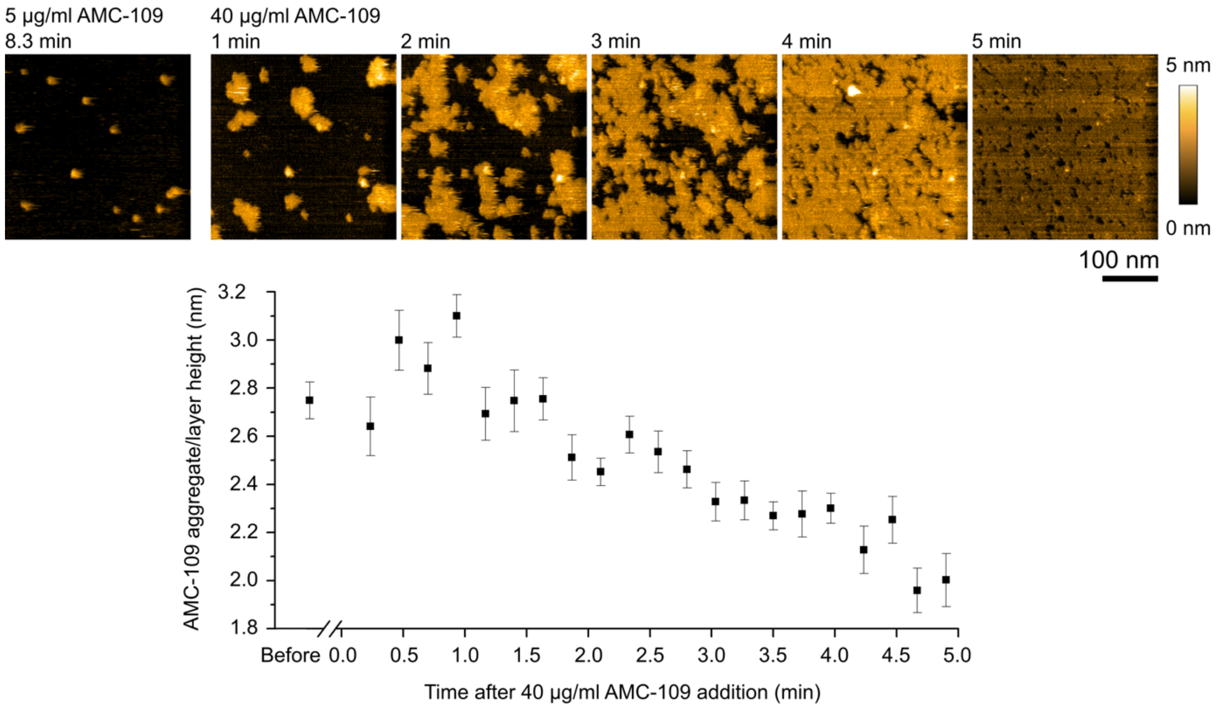


**Fig. S6**: AMC-109 on the mica background in HS-AFM imaging. In order to perform accurate thickness measurements of the lipid bilayer after addition of AMC-109 (Figure 2c), it is essential that the mica background does not get covered with AMC-109, as the membrane height is measured with respect to the mica height. Indeed, this is not the case for the used concentrations in Figure 2, as shown here: (Top left image) At 5 µg/ml AMC-109 individual aggregates attach on mica. The mica background can clearly be distinguished. (Top right images) Upon increasing the concentration of AMC-109 in the buffer to 40 µg/ml, more aggregates attach, gradually forming a uniform carpet-like layer. (Bottom) The height of the aggregates growing into the carpet-like layer decreases from 2.75 ± 0.08 nm (N = 104) for individual aggregates to 2.0 ± 0.1 nm (N = 25) at 294 s at a concentration of 40 µg/ml.

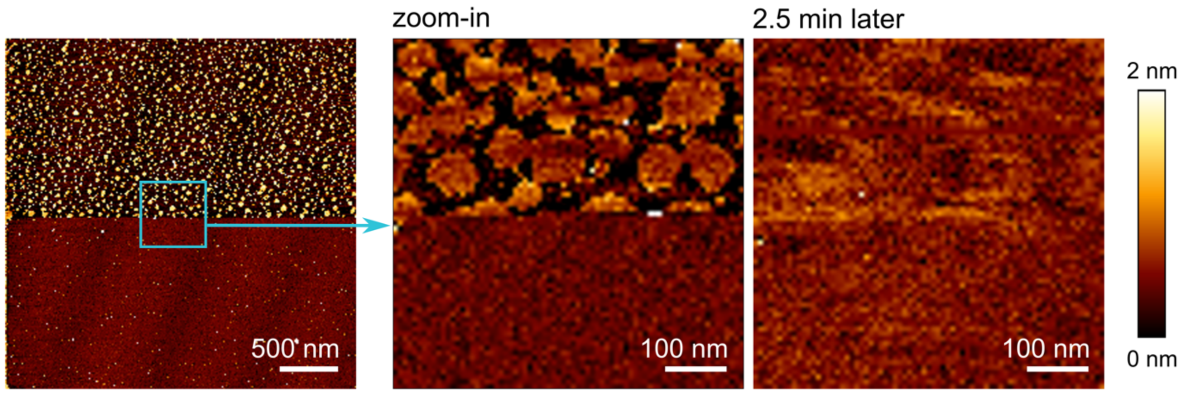


**Fig. S7**: AMC-109 on the mica background in AFM imaging. While during HS-AFM imaging no carpet-like layer of AMC-109 on the mica surface was formed during experiments at AMC-109 concentrations ≤ 5 µg/ml (Figure 2

and Figure S6), this is the case for AFM imaging. In particular in figure 1b it can be seen that the background height on the mica gradually increases after adding increasing amounts of AMC-109. A scratching experiment confirmed this. (Left) AFM imaging of a sample where 2 µg/ml AMC-109 was added to the PBS buffer onto a clean mica surface. The surface in the upper half of the left image was scratched in contact mode with 3 nN force. Next an image was taken of the scratched area (top) with the surrounding untouched layer (bottom) by AFM imaging in QI mode with 80 pN imaging force. The scratched material formed individual aggregates (bright yellow) attached on mica (black). Subsequent zoom-in image (middle) shows that the aggregates collapse again and reform the homogenous carpet-like layer. (Right) The carpet-like layer is restored within a few minutes. The height of the AMC-109 layer was estimated from the cross-sections through the AMC-109 layer and the adjacent mica surface as ~1 nm. The difference with HS-AFM imaging, where no carpet-like structure was observed for similar concentrations (Figure S6), is striking and is likely due to the longer experimental times and higher imaging force used at the AFM setup.

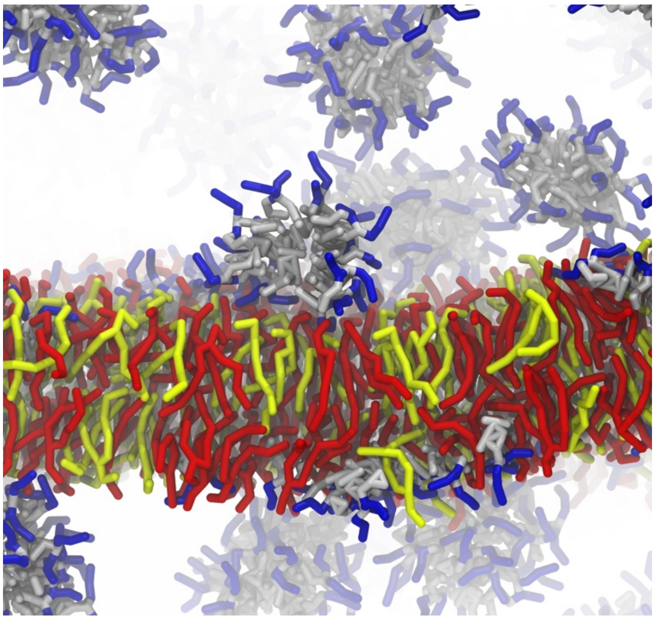

**Fig. S8**: Snapshot of the simulation box showing interaction of AMC-109 with the phospholipid membrane. Here, membrane consists of 50% POPC (yellow), 50% POPG (red). Multiple aggregates of AMC-109 (blue-grey) spontaneously formed in the bulk of water (white) and started attaching to the membrane from both sides.

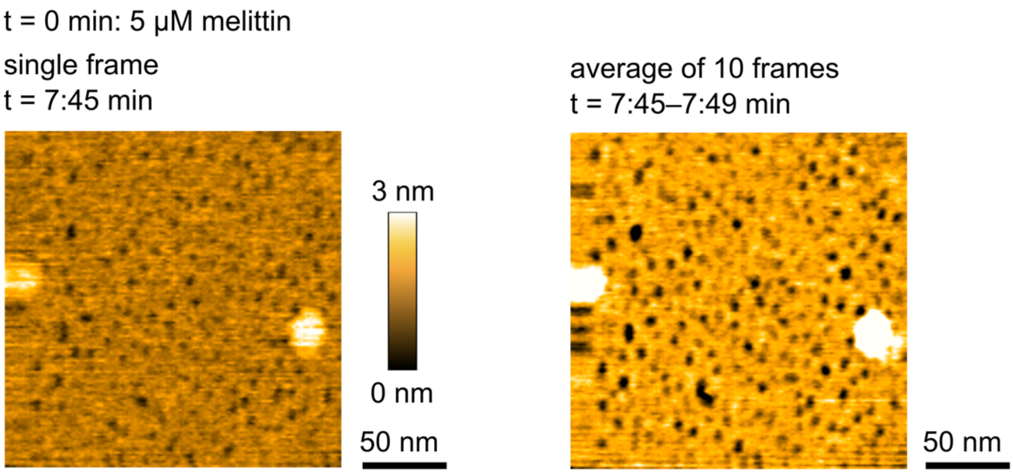


**Fig. S9**: Melittin induced pores in *S. aureus* lipid membrane visualized by HS-AFM after the addition of 5 µM melittin. (Left) Single frame from the HS-AFM video with the time step of 0.5 s. (Right) Superposition of 10 consecutive frames showing average intensity for each pixel from the superimposed frames. Individual pores in the membrane are visible as black dots and lateral domains as bright circles.

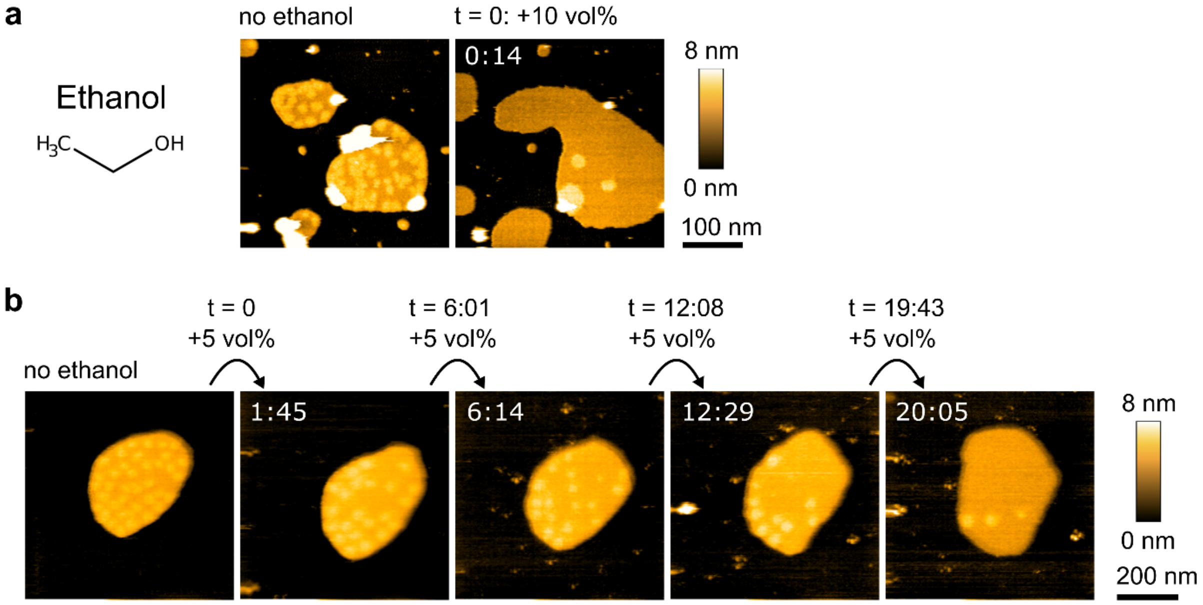


**Fig. S10**: Effects of ethanol on *S. aureus* lipid membranes observed by HS-AFM. **a** Chemical structure of ethanol and the membrane before and after the addition of 10 vol% of ethanol. We observe immediate domain dissolution and slight stretching of the membrane. **b** *S. aureus* lipid membrane before and after step by step addition of 5 vol% of ethanol. In order to observe the changes in more detail than in panel **a**, we decreased the amount of ethanol added in each step for this experiment. Some of the domains are dissolved within a few seconds after each addition of 5 vol% of ethanol. Slight stretching of the membrane is also observed.

### References used in Supporting Information:
